# Senescent myoblasts exhibit ROS-dependent Akt-mTORC1 dysregulation and are susceptible to reductive stress-induced cell death

**DOI:** 10.64898/2026.04.12.717935

**Authors:** Vladimir Belhac, Alex Dillingham, Emilia Coward, Bradley Teal, Mark C. Turner, Stephanie D. Gagnon, Jiani Qian, Harry Wilford, Ella Warren, Natalie Moger, Bernadette Carroll, Owen G. Davies, Hannah F. Dugdale, Neil R.W. Martin

**Affiliations:** School of Sport, Exercise and Health Sciences, Loughborough University, Loughborough, UK; Centre for Biological Engineering, Wolfson School of Mechanical, Electrical and Manufacturing Engineering, Loughborough University, Loughborough, UK; Research Centre for Health and Life Sciences, Coventry University, Coventry, UK; Institute for Cardio-Metabolic Medicine, University Hospitals Coventry and Warwickshire, Coventry, UK; Perelman Centre for Cellular and Molecular Therapeutics, Children’s Hospital of Philadelphia, Pennsylvania, USA; Randall Centre for Cell and Molecular Biophysics, King’s College London, Guy’s Campus, London, UK; School of Biochemistry, University of Bristol, Bristol, UK

**Keywords:** Protein synthesis, oxidants, ageing, senotherapeutic, skeletal muscle

## Abstract

Ageing is characterised by the accumulation of senescent cells. Owing to their irreversible cell-cycle arrest, these cells lack the capacity to replenish the stem cell pool and regenerate tissue, while their pro-inflammatory secretome propagates senescence in a paracrine manner. Much of the senescent phenotype has been attributed to dysregulated mTORC1 signalling, a key regulator of protein synthesis implicated in organismal ageing. Nonetheless, the mechanism underlying this dysregulation is poorly understood and limited to a few selected cell types. Here, we show that mTORC1 dysregulation is also a characteristic of senescent muscle precursor cells, and in contrast to reports in other cell types, senescent myoblasts do not rely on lysosomal nutrient liberation to sustain mTORC1 activity. Instead, they appear to depend on the PI3K/Akt pathway, which is upregulated in these cells. Exogenous antioxidants were identified to alleviate PI3K/Akt/mTORC1 signalling, while exogenous ROS has the capacity to activate mTORC1, supporting a model in which ROS acts upstream of this pathway in senescent myoblasts. Moreover, antioxidants were able to suppress the expression of pro-inflammatory cytokines and enhance the differentiation of senescent myoblasts. Interestingly, prolonged antioxidant treatment led to increased cell death in senescent but not proliferating myoblasts, suggesting they are more prone to reductive stress-induced cell death. We propose that, *in vitro*, the antioxidant capacity of many plant-derived compounds may underlie their reported benefits as therapeutics targeting senescent cells (senotherapeutics). Together, our findings provide novel insights into mTORC1-dependent regulation of the senescent phenotype and highlight the role of redox modulation in senotherapeutic strategies.

## Introduction

The mechanistic Target Of Rapamycin Complex 1 (mTORC1) is a key regulator of cellular metabolism and acts to coordinate nutrient status with cellular growth (Liu & Sabatini 2020). Specifically, amino acid and growth factor availability activate mTORC1 through a coincidence detector mechanism, resulting in phosphorylation of downstream effectors which enhance macromolecule synthesis, and decrease macromolecule breakdown at the lysosome via autophagy. Ageing has consistently been associated with dysregulated mTORC1 signalling in several tissues across multiple species (Hua et al. 2011; Kim et al. 2020; Sengupta et al. 2010; Tang et al. 2019), and its pharmaceutical inhibition enhances mammalian lifespan (Harrison et al. 2009; Komarova et al. 2012), delays tumorigenesis (Komarova et al. 2012), and counteracts age-related muscle loss (Joseph et al. 2019).

Dysregulated mTORC1 signalling is also a feature of cellular senescence (Carroll et al. 2017), a state of irreversible cell cycle arrest induced by replicative exhaustion or cellular stresses (Di Micco et al. 2021). Senescent cells accumulate with ageing, contributing to tissue dysfunction. For instance, myogenic stem cells can become senescent in ageing, hindering their capacity to regenerate skeletal muscle fibres upon injury (Sousa-Victor et al. 2014). In addition to their impaired function, senescent cells contribute to organismal ageing in a paracrine fashion via a senescence-associated secretory phenotype (SASP). Unsurprisingly, mTORC1 has been shown to underlie several senescent phenotypes, including SASP (Herranz et al. 2015; Laberge et al. 2015) and mitochondrial dysfunction (Correia-Melo et al. 2016).

Despite the essential role of mTORC1 dysregulation in both the senescent cell phenotype and organismal ageing, the mechanisms underlying this dysregulation remain largely unexplored and have been studied primarily in fibroblasts. Here, we investigated mTORC1 signalling in myoblasts, committed myogenic progenitor cells, which possess the capacity to differentiate and fuse into a mature muscle fibre (Smith et al. 2023). Evidence suggests that senescence of myogenic cells contributes to skeletal muscle ageing (Sousa-Victor et al. 2014), which is characterised by reduced muscle mass, strength, and quality of life, resulting in financial burden on health and social care services (Pinedo-Villanueva et al. 2019). Therefore, studying mTORC1 in senescent myogenic cells may not only advance our understanding of its dysregulation in senescence but also helps identify strategies to alleviate aspects of skeletal muscle ageing and improve quality of life.

Together, the objectives of the present experiments were threefold. Firstly, to determine whether senescent committed muscle progenitor cells exhibit dysregulated mTORC1 signalling. Secondly, to identify the upstream signals responsible for driving this dysregulation. Finally, to test whether alleviating mTORC1 dysregulation can reduce or reverse key features of the senescent phenotype. Using C2C12 and primary human myoblasts, we show herein that senescence is associated with chronically activated mTORC1 that is desensitised to nutrient starvation. This dysregulated mTORC1 signalling could not be restored by inhibition of autophagy but instead appeared to be regulated through Akt (growth factor signalling) and mediated by reactive oxygen species (ROS). Finally, we demonstrate that chronic treatment with antioxidants can selectively eliminate senescent, but not proliferating cells, leading us to postulate that senescent cells are more susceptible to reductive stress induced cell death. These data provide novel insights into the biology of skeletal muscle ageing and may contribute to the development of therapeutics to alleviate age-related muscle loss.

## Results

### mTORC1 is dysregulated in senescent myoblasts

Our initial experiments sought to induce stress-induced senescence in C2C12 myoblasts. We found that incubating cells with 5 µM of the DNA topoisomerase inhibitor, etoposide, increased cell size, the proportion of β-galactosidase-positive cells and elevated mRNA expression of senescence-associated genes (Figure 1A-C), all indicative of a senescent phenotype.

**Figure 1.**
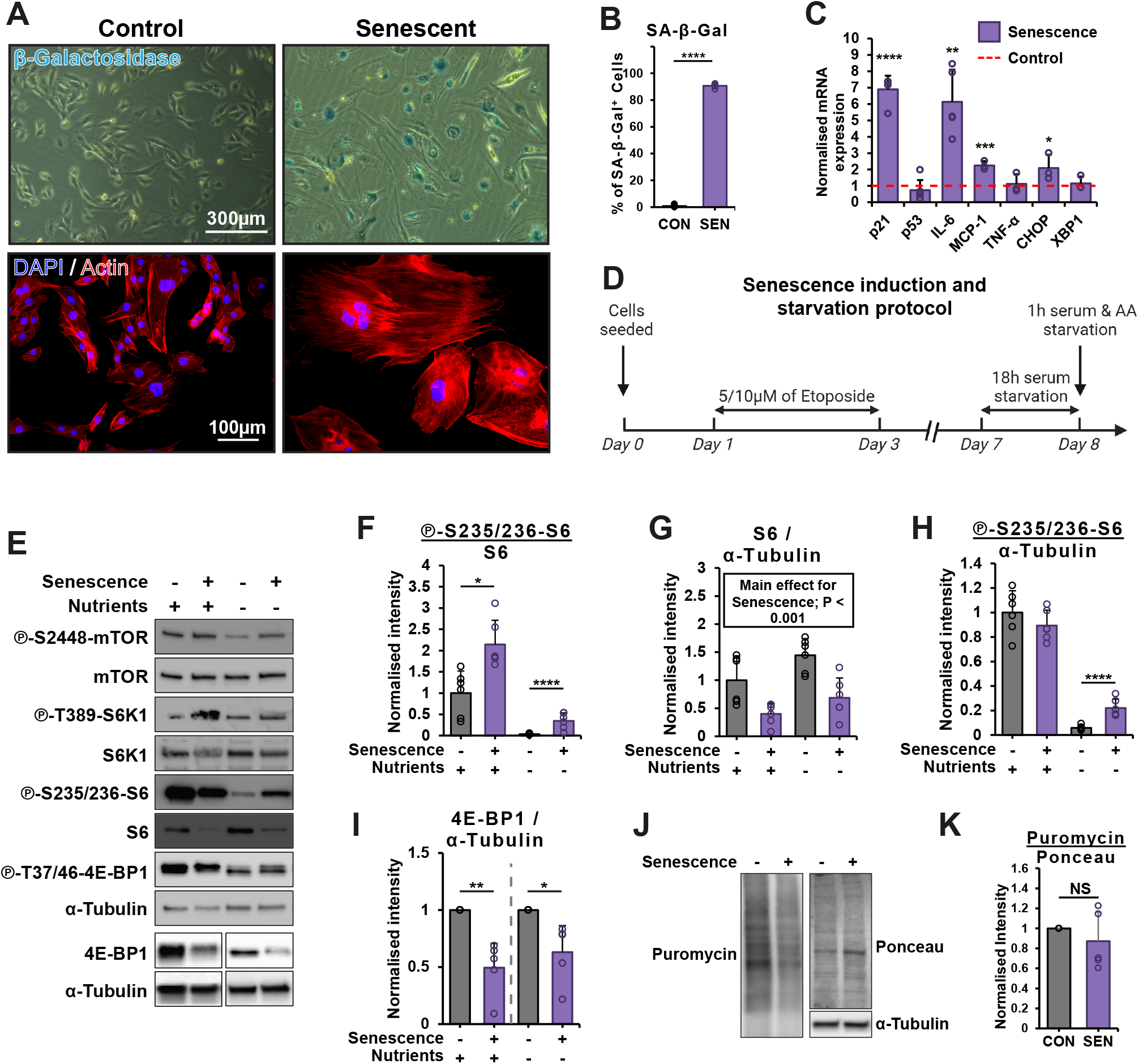
Senescent myoblasts exhibit dysregulated mTORC1 signalling. C2C12 myoblasts were treated with 5 µM etoposide for 48 h and then maintained in growth medium for an additional five days. (A and B) Images of cells stained for F-actin to assess morphology and senescence-associated β-galactosidase to assess lysosomal content, with quantification for the latter shown in (B). (C) mRNA expression of senescence-associated genes quantified (D) Schematic summarising the senescence induction and starvation protocol. Cells were starved by 18 h serum deprivation followed by 1 h combined serum and amino acid deprivation. E-K) Western blot analysis and quantification of mTORC1 signalling (E-I) and puromycin incorporation to assess protein synthesis (J and K) in C2C12 myoblasts. Results are reported as mean ± standard deviation. P values were calculated using independent, heteroscedastic t-test (B, C, and K) or independent Two-way ANOVA followed by followed by calculating estimated marginal means with Bonferroni correction (F-I). n = 3-6. *P ≤ 0.05, **P ≤ 0.01, ***P ≤ 0.001. CON = control, SEN = senescent.

To investigate mTORC1 signalling in senescent myoblasts, we considered previous research suggesting that mTORC1 dysregulation is only detectable after the manipulation of its key activators, nutrients and growth factors (Carroll et al. 2017). Therefore, we assessed mTORC1 in full GM (Growth Media) and serum- and amino acid-starved conditions (Figure 1D for protocol summary). The results revealed that, regardless of nutrient availability, senescent myoblasts displayed elevated phosphorylation of all measured proteins downstream of mTORC1, including S6K1, S6, and 4E-BP1 (Figure 1E and F). However, the total abundance of these same targets was reduced in senescent myoblasts (Figures 1E, G and I), hence the results were also normalised for housekeeping protein (α-Tubulin), where relevant, so that we could better assess the anabolic effect downstream of mTORC1. Once normalised for a housekeeping protein, some mTORC1 downstream targets, namely S6 and 4E-BP1, were elevated only in the absence of nutrients (Figure 1H). This reinforced the idea that mTORC1 dysregulation or its downstream signalling is most evident during nutrient starvation, and thus we studied mTORC1 dysregulation primarily in the starved state hereafter. Additionally, total protein synthesis measured by puromycin was not significantly altered in senescent myoblasts (Figure 1J and K).

We further proceeded to validate our findings in primary human myoblasts, demonstrating that treatment with 10 µM etoposide increased the proportion of β-galactosidase-positive cells, along with enhanced phosphorylation of mTORC1-associated residues during nutrient starvation (Figure S1). Collectively, these results suggest that senescent myoblasts exhibit dysregulated mTORC1 signalling.

### mTORC1 dysregulation is dependent on PI3K/Akt and not autophagy

To investigate the mechanism underlying mTORC1 dysregulation, we assessed its canonical regulators: the nutrient-sensing arm, which facilitates mTOR translocation to the lysosome, and the insulin/growth factor signalling arm, which facilitates its kinase activation at the lysosome via the PI3K/Akt signalling pathway (See Figure 2A). Existing evidence from non-myogenic cells indicates that senescence is characterised by cytoplasmic regions where mTOR and lysosomes are enriched and highly colocalized, highlighting an altered pattern of mTORC1 lysosomal translocation within the nutrient sensing arm (Narita et al., 2011). In contrast, we did not observe such a pattern in C2C12 myoblasts, nor did we detect any absolute difference in mTOR colocalization with lysosomes, as measured by its colocalization with the lysosomal protein LAMP1 (Figures 2B and C).

**Figure 2.**
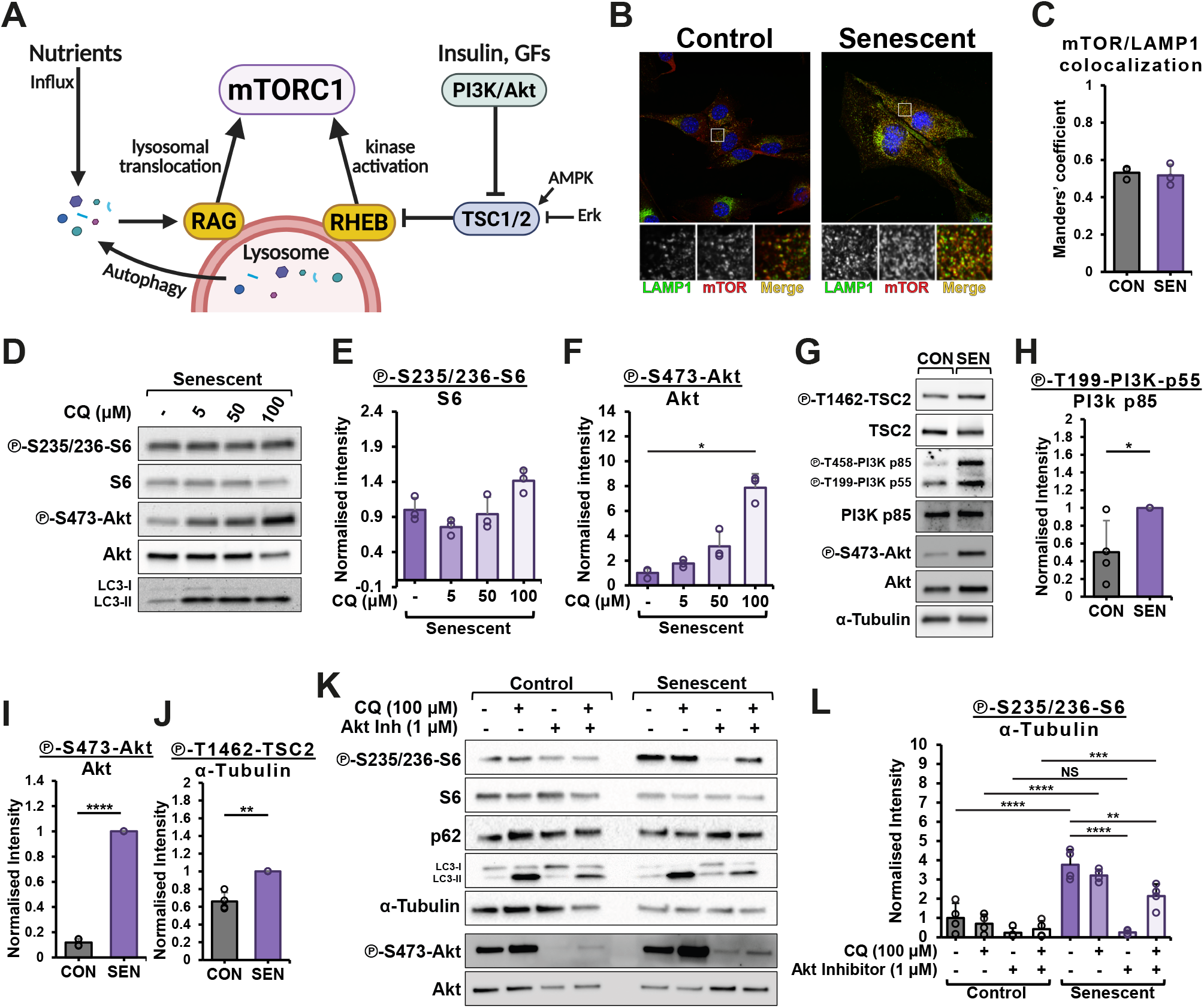
Inhibition of Akt, but not autophagy, alleviates mTORC1 activity in starved senescent C2C12 myoblasts. (A) mTORC1 regulation. The presence of nutrients, particularly essential amino acids, either through extracellular influx or intracellular regeneration, facilitates the translocation of mTORC1 to the lysosomal surface via Rag GTPases. Once localized to the lysosome, mTORC1 kinase activity is subsequently activated through the PI3K/Akt/TSC2/Rheb GTPase signalling pathway. (B and C) Immunostaining for mTOR and LAMP1 in control and senescent myoblasts (B) and colocalization analysis using Mander’s coefficient (C). (D-L) Western blot analysis and quantification for mTORC1 and PI3K/Akt/TSC2 signalling in control and senescent myoblasts treated with either vehicle, CQ, or Akt Inh as indicated. P values were calculated using an independent, heteroscedastic t-test (C, H-J), Kruskal-Wallis test followed by Dunn’s test (F), and two-way ANOVA followed by estimated marginal means with Bonferroni correction (L). n = 3–4. *P ≤ 0.05, **P ≤ 0.01, ***P ≤ 0.001, ****P ≤ 0.0001. CQ = chloroquine, Akt Inh = Akt inhibitor, CON = control, SEN = senescent.

However, senescent myoblasts exhibited increased lysosomal content, measured by β-galactosidase assay (Figures 1A and B) and immunoblotting for LAMP1 (Figure S2A), as well as elevated lysosomal pH (Figure S2B, see Supplementary Methods), in line with previous reports suggesting that increased lysosomal biogenesis may compensate for lysosomal dysfunction (Curnock et al. 2023). Further evidence from non-myogenic cells indicated that mTORC1 dysregulation can be dependent on autophagy (Carroll et al. 2017; Jiang et al. 2022), a catabolic process in which cellular components are degraded inside lysosomes, releasing nutrients capable of activating mTORC1 (Yu et al. 2010). Nonetheless, autophagic flux, measured by LC3-II turnover, was not significantly different in senescent myoblasts (Figure S2C-E). To determine whether autophagy inhibition alleviates mTORC1 signalling, senescent myoblasts were treated with chloroquine (CQ) or bafilomycin A during the final hour of starvation. These treatments did not reduce the phosphorylation of the mTORC1 downstream effector S6 and, in some cases, even increased it (Figure 2D-E and Figure S2F), suggesting that autophagy inhibition does not suppress mTORC1 activity.

CQ also dose-dependently upregulated Akt phosphorylation (Figure 2F), raising the question of whether Akt and insulin/growth factor signalling sustains mTORC1 when autophagy is inhibited. This led us to shift our focus to the second arm of mTORC1 regulation, involving its kinase activation, in which Akt is a key intermediate (Figure 2A). Senescent myoblasts showed substantially elevated phosphorylation of PI3K, Akt, and TSC2 (Figure 2G and J). Irrespective of concentration, the Akt catalytic inhibitor significantly reduced S6 phosphorylation by ∼90%, indicative of decreased mTORC1 signalling (Figure S2G and H). Moreover, the S6 phosphorylation levels became almost equal between Akt inhibited control and senescent cells (Figures K and L), suggesting that mTORC1 dysregulation in senescent myoblasts is primarily driven by its increased kinase activation. Interestingly, we also found that the combination of Akt inhibitor and CQ antagonise each other, as Akt, S6 and LC3-II are closer to baseline than when each inhibitor is administered alone (Figure 2K).

### ROS modulate PI3K/Akt/mTORC1 signalling

A hallmark of senescent cells is the production of a secretome termed the SASP (senescence-associated secretory phenotype), which is rich in growth factors, cytokines, and ROS (Di Micco et al. 2021), which act upstream of PI3K/Akt signalling (Sies & Jones 2020; Thorpe et al. 2015). A series of experiments were conducted, in which the components of SASP were targeted to test whether they act upstream of Akt/mTORC1 dysregulation. For example, inhibiting cytokine-mediated signalling using a catalytic JAK inhibitor, Tofacitinib, did not affect Akt/mTORC1 (Figure S3A-C). However, the antioxidant Tiron inhibited both Akt and mTORC1 phosphorylation by up to a half, suggesting that ROS may mediate the dysregulation of this signalling axis. By scavenging superoxide, Tiron can elevate cellular levels of H_2_O_2_ (Matoba et al. 2003), which is involved in PI3K/Akt signalling (Sies & Jones 2020). Therefore, we further validated our results using NAC (N-acetylcysteine), which is more effective at scavenging H_2_O_2_, partly through synthesis of glutathione, a central component of intracellular antioxidant defence. Similar to the above, 1 h of NAC treatment significantly alleviated Akt/mTORC1 signalling in nutrient starved senescent myoblasts (Figure S3D and E) and remained inhibited when the treatment was extended to 19 hours (>60% reduction for Akt and S6 phosphorylation; Figures 3A-C). Redox-sensitive AMPK phosphorylation at T172, which inhibits mTORC1, was unchanged in response to NAC, suggesting that these changes happen independent of AMPK (Figures S3F). Interestingly, 1 h of NAC in senescent cells in full GM dose-dependently alleviated Akt phosphorylation, but only a high (20 mM) dose inhibited S6 phosphorylation (Figures S3G-I).

**Figure 3.**
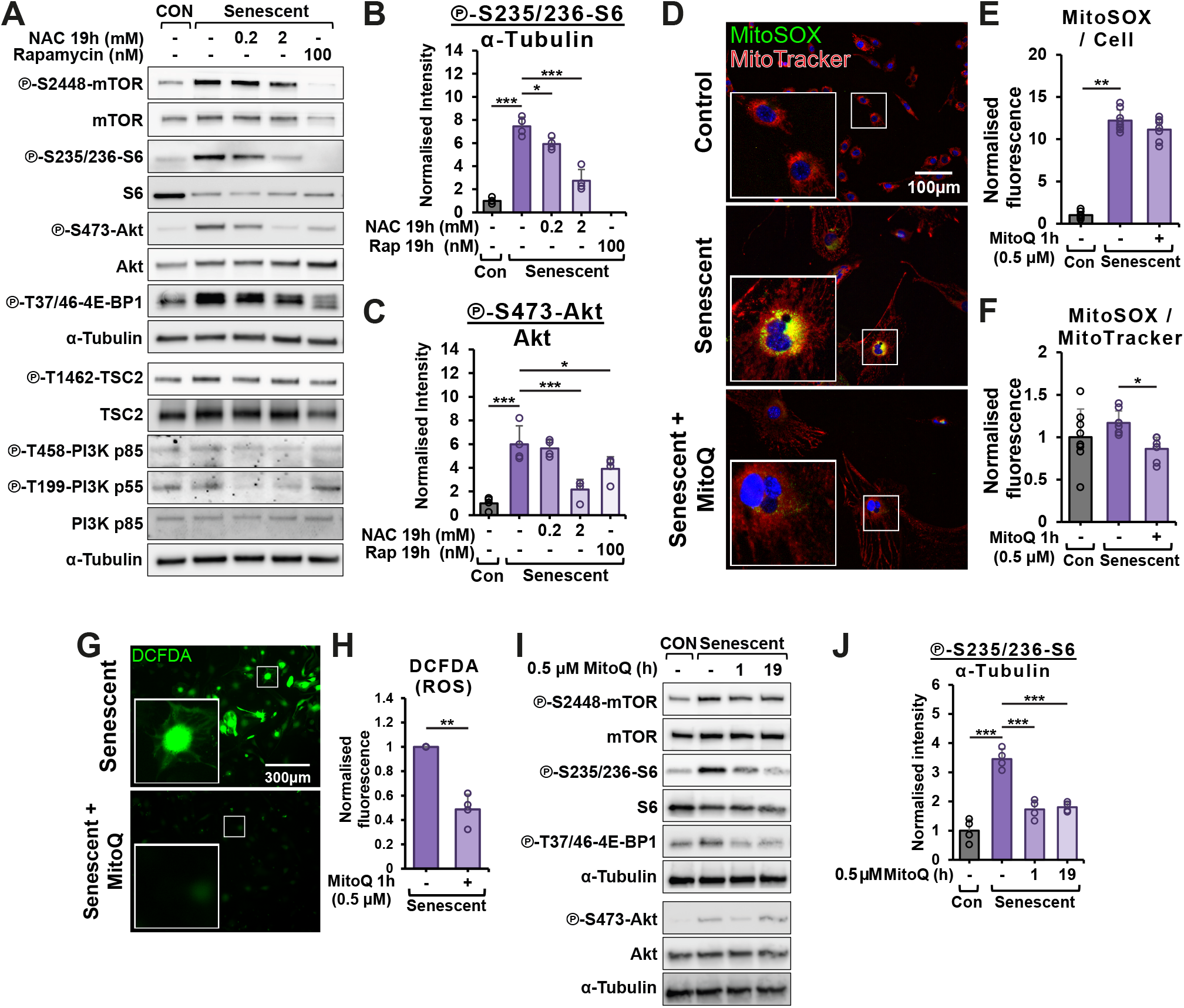
ROS contributes to Akt/mTORC1 dysregulation in senescent C2C12 myoblasts. (A–C) Western blot analysis of Akt/mTORC1 signalling in starved myoblasts treated with the indicated doses of NAC. (D–F) Fluorescence microscopy images and quantification of MitoSOX Red and MitoTracker Deep Red staining intensity in control and senescent cells, with or without MitoQ. (G and H) Fluorescence microscopy images and quantification of DCFDA ROS staining in senescent myoblasts with or without MitoQ. (I and J) Western blot analysis of Akt/mTORC1 signalling in starved control and senescent cells treated with 0.5 µM of MitoQ for the indicated time. P values were calculated using an independent, heteroscedastic t-test (H), and one-way ANOVA followed by Dunnett’s post hoc test (B, C, E, F, J). n = 4–8. *P ≤ 0.05, **P ≤ 0.01, ***P ≤ 0.001. NAC = N-acetylcysteine, MitoQ = Mitoquinone mesylate, CON = control, SEN = senescent.

Since antioxidants had profound effects on Akt-mTORC1 signalling, next we characterised ROS production and related gene expression in senescent myoblasts. Immunoblotting analysis revealed a decrease in both total and phosphorylated Nrf2 (Figures S3J and K), a master regulator of the cellular antioxidant response, yet unexpectedly showed increased expression of the downstream endogenous antioxidants SOD2 and PRDX6 (Figure S3J). Furthermore, we found an increased expression of NADPH oxidase 4 (Figure S3J), a key source of ROS and the most abundant NOX isoform in C2C12 myoblasts (Handayaningsih et al. 2011). We then used the DCFDA ROS assay, sensitive to H_2_O_2_-derived radicals, peroxynitrite, and other ROS, and found that senescent myoblasts generate more ROS, while NAC effectively reduces it (Figure S3L). Senescent myoblasts further exhibited increased MitoSOX staining, indicative of increased mitochondrial superoxide production (Figures 3D and E). However, this difference was not statistically significant when normalised for total mitochondria (*P*=0.312), quantified through MitoTracker staining (Figures 3D and F). This reflects an increase in mitochondrial content measured by MitoTracker staining and immunoblotting for OXPHOS complexes (Figures S3M and N), which together suggest increased mitochondrial content both per cell and at the total protein level, as previously reported in senescent cells (Correia-Melo et al. 2016). Given this fact, we speculated that although the amount of ROS released per mitochondria might be the same, the increased mitochondrial content causes an increase in net ROS production per cell and total protein. Therefore, mitochondrial ROS production in senescent myoblasts was targeted by a one-hour treatment with MitoQ (mitoquinone mesylate), a ubiquinone conjugated to a lipophilic cation that enables its accumulation within mitochondria, where it scavenges ROS (Kelso et al. 2001). MitoQ significantly decreased MitoSOX (Figures 3D-F), DCFDA ROS staining (Figures 3G and H), and Akt/mTORC1 activity (Figures 3I and J), albeit Akt phosphorylation returns to baseline over time during chronic (19 h) treatment (Figure 3I).

If ROS acts upstream of Akt/mTORC1 dysregulation in senescence, increasing ROS in non-senescent cells should dysregulate Akt/mTORC1. Therefore, control C2C12 myoblasts were treated with H_2_O_2_ using the standard starvation protocol, with the treatment shortened to 10 minutes due to H_2_O_2_’s short half-life in cultured cells (Seaver & Imlay 2001; Gülden et al. 2010). H_2_O_2_upregulated phosphorylation of mTORC1’s downstream targets, but without significantly affecting TSC2 and Akt phosphorylation (Figures S3O-R), suggesting that mTORC1 can also be regulated by ROS in a manner that is independent of PI3K and Akt.

### Antioxidants improve differentiation of senescent myoblasts

MTORC1 signalling has been previously described as a key mediator of phenotypes characteristic of senescent cells. This includes the synthesis of SASP through increased selective protein translation and indirectly increasing their mRNA expression (Herranz et al., 2015; Laberge et al., 2015). One would predict that if antioxidants can inhibit these pathways, they would in turn also downregulate the production of SASP. Indeed, we found that NAC decreased the mRNA expression of cytokines IL-6 and MCP-1 (Figure 4A and B).

**Figure 4.**
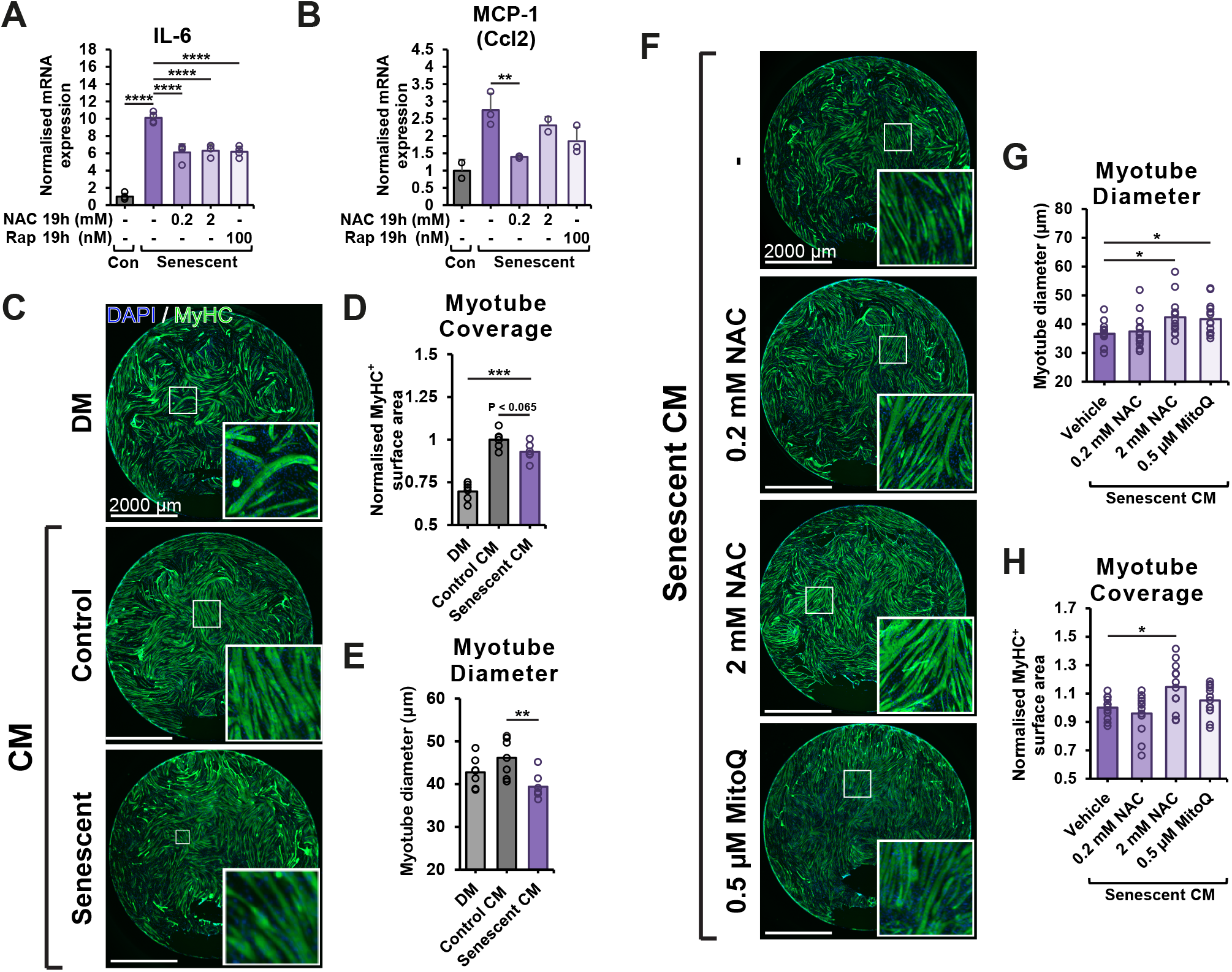
Antioxidants alleviate cytokine mRNA expression and improve differentiation of C2C12 myoblasts. (A, B) C2C12 myoblasts were starved following the standard protocol and treated with indicated doses of NAC as in Figure 3A. mRNA expression of indicated genes was measured using qPCR. (D–F) Representative images and quantification of myoblast differentiation in a 96-well plate after treatment with conditioned medium collected from control or senescent myoblasts. (G-I) As in (D–F), but prior to differentiation, cells were pretreated with antioxidants for 24 h, as indicated. Results are reported as mean ± standard deviation. P values were calculated using one way ANOVA followed by Dunnett’s post hoc test. n = 3-14. *P ≤ 0.05, **P ≤ 0.01, ***P ≤ 0.001, ****P≤ 0.0001. NAC = N-acetylcysteine, MitoQ = Mitoquinone mesylate, CON = control, SEN = senescent.

SASP is also reported to diminish cell differentiation (Baar et al. 2018). To test the same conclusions in our model of myoblast senescence, we differentiated C2C12 myoblasts using conditioned medium, prepared from differentiating control and senescent myoblasts (protocol summary in Fig. 4SA). Conditioned media from differentiating senescent myoblasts caused a small decrease in myotube (MyHC^+^) thickness and surface coverage of C2C12 myoblasts (Figure 4C–E), highlighting their decreased differentiation capacity. Interestingly, pre-treating senescent cells with antioxidants alleviated this effect of conditioned media by 10-20% (Figure 4F–H). Additionally, we found that senescent myoblasts almost completely lose their capacity to differentiate, and antioxidant treatment caused a marginal but statistically significant increase in myotube coverage (Figure S4B and C). The latter results are also unlikely to be the consequence of improved cell survival commonly observed with antioxidants, given that antioxidants did not result in higher cell count, measured by DAPI staining (Figure S4B and D).

### Senescent myoblasts are more susceptible to reductive stress-induced cell death

Senescent cells tend to exhibit resistance to apoptosis, facilitated by altered cell survival and death signalling (Di Micco et al. 2021), a finding we also observed in senescent C2C12 myoblasts (Figures S5A and B), with BAD phosphorylation downstream of Akt also being redox sensitive (Figures S5C-F). Despite this, we repeatedly observed that senescent myoblasts treated chronically with antioxidants show signs of cell stress/cytotoxicity. For instance, in experiments where senescent cells were treated with antioxidants for 19 hours during nutrient starvation, a higher NAC dose (2 mM) led to increased mRNA expression of genes associated with DNA damage and ER stress (Figure S5G-I). Furthermore, prolonged antioxidant treatment during myoblast differentiation in low-serum medium resulted in reduced cell count (Figure 5A derived from Figures S4B and D) and protein content (Figure 5C). We subsequently confirmed increased cleaved caspase-3 expression, indicating enhanced cell death (Figures 5B and D). In addition, the DCFDA ROS assay showed that prolonged MitoQ treatment failed to reduce total ROS levels (Figure 5E and F), in contrast to an acute treatment (Figure 3G and H). Interestingly, MitoQ treatment markedly increased cell-to-cell variability in DCFDA staining intensity (Figure 5G), indicating that more cells are residing at the ends of the oxidative– reductive stress continuum.

**Figure 5.**
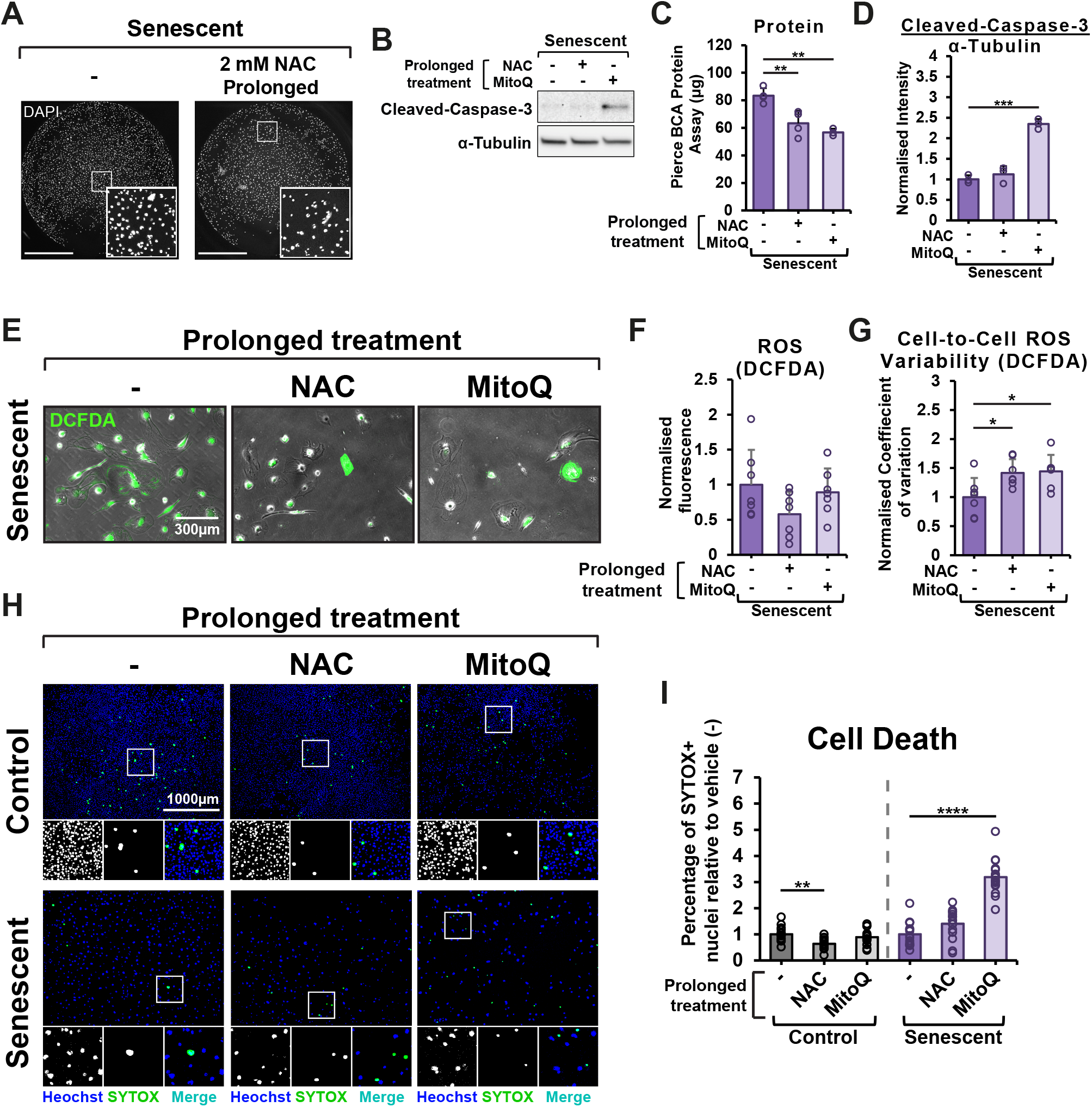
Senescent C2C12 myoblasts are susceptible to antioxidant-induced cell death. A-G) Senescent myoblasts, exposed to five days of differentiation protocol and prolonged treatment with a vehicle (-), 2 mM of NAC, or 0.5 µM of MitoQ, before immunostaining (A; image extracted from Figure S4B) or protein concentration quantification (BCA protein assay; C) and Western blotting for cleaved caspase expression (B and D), as well as being stained and imaged for ROS using the DCFDA assay (E-G). (H and I) Live and dead cell staining using Hoechst 33342 and SYTOX, respectively, of myoblasts treated with a vehicle (−), 2 mM of NAC, or 0.5 µM of MitoQ for three days, with the first 19 h of treatment being exposed to the standard starvation protocol, composed of 18 h serum deprivation, followed by 1 h of both serum and amino acids deprivation. Results are reported as mean ± standard deviation. P values were calculated using one-way ANOVA followed by Dunnett’s post hoc test. n = 3-16. *P ≤ 0.05, **P ≤ 0.01, ***P ≤ 0.001, ****P ≤ 0.0001. NAC = N-acetylcysteine, MitoQ = Mitoquinone mesylate, CON = control, SEN = senescent.

Since any compound can become cytotoxic at a sufficient concentration or in the right context, we next tested whether the antioxidant-mediated cytotoxicity is specific for senescent cells. This was largely inspired by the similarities between our results with antioxidants and previously reported compounds with the capacity to selectively induce cell death in senescent cells, termed senolytics. Some senolytics have potent antioxidant properties and the proposed mechanism underlying their senolysis is the inhibition of the PI3K/Akt/mTORC1 cascade (Di Micco et al. 2021), which our data suggest is inhibited by antioxidants alone. To test this idea directly, we performed SYTOX staining for cell viability, which revealed that antioxidants decreased the proportion of healthy myoblasts with permeabilised nuclei but increased it in senescent myoblasts (Figure 5H and I), confirming that antioxidants can induce cell death selectively in senescent cells.

## Discussion

Accumulation of senescent cells contributes to ageing phenotypes and tissue dysfunction (Di Micco et al. 2021). Because senescent cells do not proliferate, one would suspect that the anabolic signalling cascades, such as mTORC1, would be inhibited, a fate observed in quiescent cells (Rodgers et al. 2014). However, the existing evidence suggests this is not the case, instead mTORC1 is reported to be desensitised to nutrient starvation (Carroll et al. 2017; Jiang et al. 2022) or chronically elevated relative to proliferating cells (Zhang et al. 2024; Cho et al. 2020). Our data from C2C12 myoblasts indicate a combination of these phenomena. Irrespective of nutrient availability, the fraction of phosphorylated mTORC1 targets, S6K1, S6, and 4E-BP1, was increased, indicating elevated mTORC1 activity. However, when the phosphorylation levels were normalised to a housekeeping protein (α-Tubulin), it revealed that an increase in absolute phosphorylation of S6 and 4E-BP1 occurs only under nutrient-starved condition. The mechanism underlying the nutrient-dependent effect is not entirely clear but may be linked to oversaturation of growth factors and amino acids from high serum growth media or other unknown properties of senescent cells. The discrepancy between fractional and absolute phosphorylation is mediated by the decrease in total expression of the measured downstream targets of mTORC1 in senescent cells. This might be particularly consequential for 4E-BP1, which sequesters eIF4E and inhibits cap-dependent translation. Its decreased expression in senescent cells might therefore enhance protein translation. mTORC1 regulates 4E-BP1 by inhibitory phosphorylation, which suggests 4E-BP1 in senescent myoblasts is also subject to mTORC1-independent regulation. Despite not proliferating, senescent myoblasts did not show a reduction in protein synthesis, as measured by the SUnSET assay. While mTORC1 enhances global protein translation, it also selectively promotes the translation of specific mRNAs, including those associated with SASP (Herranz et al., 2015; Laberge et al., 2015). Its increased activity, coupled with unchanged global protein synthesis, might therefore indicate a shift toward a more mTORC1-specific translatome.

In contrast to studies in other cell types (Narita et al. 2011; Carroll et al. 2017; Jiang et al. 2022), our results in myoblasts indicated that the nutrient-sensing arm, facilitating mTORC1’s translocation to the lysosome, is not affected in senescent myoblasts. While mTOR–LAMP1 colocalization was used as a key measurement supporting our conclusion, we recognise that this approach has inherent limitations, particularly those introduced by differences in cell size between control and senescent cells, including an altered likelihood of random colocalization. It should be also noted that mTOR’s lysosomal localization in myogenic cells is less well defined than in other cell types, with reported changes in mTOR–LAMP1 colocalization remaining relatively small in response to stimuli (Wang et al. 2021). Subsequent experiments showed that inhibiting autophagy, which has the capacity to generate nutrients for mTORC1 reactivation (Yu et al. 2010), does not alleviate the dysregulation of mTORC1. We suspect that this may be a consequence of a concomitant increase in Akt phosphorylation upstream of mTORC1, which we have shown in response to autophagy inhibitors such as chloroquine. Increased Akt phosphorylation has been previously reported following autophagy inhibition specifically in muscle cells (Halaby et al. 2013; Spears et al. 2016; Zhou et al. 2016; Ryan et al. 2024), but to our knowledge, not in other cell types (Zou et al. 2013), highlighting a potential cell type specific response. Furthermore, our data demonstrates that the PI3K/Akt signalling pathway, which regulates kinase activity of mTORC1, is hyperphosphorylated in nutrient starved senescent myoblasts. Inhibition of Akt inhibits mTORC1 to the levels observed in Akt inhibited control cells, suggesting this pathway mediates mTORC1 dysregulation in senescent myoblasts.

Subsequent experiments demonstrated that antioxidants alleviate the hyperphosphorylation of Akt/mTORC1 in C2C12 myoblasts. Other studies have demonstrated that enhancing antioxidant capacity *in vitro* through pharmaceutical or genetic means can prevent the acquisition of Akt or mTORC1 dysregulation, as well as the senescent phenotype in general (García-Prat et al. 2016; Summer et al. 2019; Yang et al. 2018; Zhang et al. 2015; Nacarelli et al. 2016). However, to our knowledge, our data are the first to demonstrate that an acute (1 h) antioxidant treatment in fully developed senescent cells is sufficient to inhibit aberrant PI3K/Akt/mTORC1 signalling, indicating that the chronic presence of ROS is not only mediating the acquisition of the senescence phenotype but also its maintenance. ROS are well documented to regulate receptor tyrosine kinases via oxidation of thiols at cysteine residues upstream of PI3K/Akt, a mechanism that was described as a contributing factor to uncontrolled growth and proliferation of cancer cells (Sies & Jones 2020). Akt/mTORC1 inhibition in response to enhanced ROS scavenging via dietary or genetic means, has also been reported in mice and human studies *in vivo* (Chen et al. 2014; Pal et al. 2014; Michailidis et al. 2013). Collectively, these findings indicate that ROS act upstream of the Akt/mTORC1 cascade and that antioxidants can be used to inhibit it.

Despite the clear relationship between Akt and mTORC1 in senescent myoblasts, mTORC1 regulation by ROS seemed to be Akt-independent in some instances. Notably, H_2_O_2_ in proliferating C2C12 myoblasts activated mTORC1 without changing Akt. This might be due to oxidation of TSC2, which has been previously reported to activate mTORC1 in response to the cysteine oxidant, phenylarsine oxide (Yoshida et al. 2011). It should be noted, however, that the manner in which H_2_O_2_ was used in this study may not adequately simulate the transient and localized ROS production that occurs endogenously (Sies & Jones 2020). Consistent with the context-dependent nature of this pathway, our results demonstrate that NAC inhibited mTORC1 downstream targets in a dose-dependent manner when starved but had limited effect in myoblasts maintained in full growth medium. A similar phenomenon was observed in RAW 264.7 macrophages, where NAC suppressed LPS-induced Erk and p38 phosphorylation under serum starvation (1% FBS) but enhanced phosphorylation when serum was abundant (10% FBS; Chan, Riches and White, 2001). Other studies have further indicated that ROS effects on mTORC1 vary depending on dose, treatment duration, and cell type (Li et al. 2010). Collectively, these findings underscore the importance of cellular context in determining how ROS modulates mTORC1 activity. While our data show that antioxidants can inhibit these pathways, we believe that future studies will need to consider its localized nature, monitoring additional ROS-sensitive effectors upstream of mTORC1, such as AMPK and MAPKs, and which oxidation sites are involved.

In addition to Akt/mTORC1 signalling, antioxidants alleviated IL-6 and CCL2 mRNA expression, key components of the SASP. This effect is likely both Akt/mTORC1-dependent and -independent, given the capacity of oxidants to modulate pro-inflammatory signalling independently of these pathways (Sies & Jones 2020). Antioxidant treatment prior to differentiation further rescued the impaired differentiation capacity induced by conditioned media derived from senescent C2C12 myoblasts, indicating an attenuation of the paracrine effects of the SASP on myogenic differentiation. Consistent with previous reports in primary myoblasts (Khor et al. 2016), antioxidant treatment in senescent C2C12 myoblasts also improved differentiation capacity of senescent cells themselves, albeit to a small magnitude. In contrast, inhibition of mTORC1 (Pollard et al. 2014; Park & Chen 2005) or ROS (Rajasekaran et al. 2020; Youm et al. 2019) has been reported to impair myoblast differentiation in non-senescent cells. Together, these results likely indicate that there is an optimal redox and mTORC1 activity window to maximise myoblast differentiation and that scavenging ROS can be utilised to improve myogenic differentiation of senescent cells.

We further demonstrated that prolonged antioxidant treatment can induce cell death in senescent, but not proliferating, C2C12 myoblasts. Antioxidant treatment alone is unlikely to be inherently cytotoxic to senescent cells. Supporting this, low doses or shorter treatment durations did not affect cell numbers in our experiments and have even been reported to reduce cell death in human myoblasts undergoing replicative senescence (Khor et al. 2017). Instead, we suggest that senescent cells may be more susceptible to reductive stress–mediated cell death. Reductive stress is defined as an excessive intracellular reducing state, caused by the accumulation of reducing equivalents such as NADPH, glutathione, and thioredoxins (Brewer et al. 2013), which can be triggered by antioxidant treatment that is prolonged or administered at high concentrations. We propose that some senescent cells are more susceptible to reductive stress or its downstream effects by the altered regulation of redox homeostasis. Although characterised by increased ROS and decreased phosphorylation and expression of a key antioxidant defence regulator, Nrf2, senescent myoblasts also exhibited increased expression in some endogenous antioxidants, namely SOD2 and PRDX6. Similarly, increased antioxidant activity (Khor et al. 2017; Borlon et al. 2007), as well as endoplasmic reticulum-specific reductive stress (Qiao et al. 2022), has been previously reported in senescent cells, suggesting that senescent cells can exhibit hallmarks of both oxidative and reductive stress depending on the context and subcellular location. Furthermore, some evidence suggests that reductive stress can be oxidative in nature. For example, an increase in reductive potential using glutathione overexpression or NAC treatment was reported to increase the oxidation of mitochondrial proteins (Zhang et al. 2012). We also observed that although acute antioxidant treatment reduced DCFDA ROS staining, the effect was not lost after prolonged treatment. Moreover, prolonged treatment increased cell-to-cell variation, suggesting a greater proportion of cells at the extreme ends of the reductive/oxidative stress distribution. In future work, it would be interesting to investigate whether cell death in senescent cells is indeed associated with localized oxidative stress.

An alternative or complementary explanation for the increased cell death observed in senescent cells is that antioxidants inhibit the pro-survival signalling downstream of the PI3K/Akt/mTORC1 axis, a pathway that senescent cells are thought rely on more heavily for survival (Di Micco et al. 2021). This dependency is exploited by senolytics, drugs that selectively eliminate senescent cells by targeting such pro-survival pathways while sparing proliferating cells. Interestingly, several senolytics also exhibit antioxidant properties. These include plant-derived compounds such as reishi and green tea extracts (Imb et al. 2024), with the most well-studied being the flavonoids fisetin and quercetin, both of which have been shown to reduce senescent cell burden *in vitro* and *in vivo* (Di Micco et al. 2021). However, their antioxidant capacity is not currently recognised as the primary mechanism of action. Instead, inhibition of key signalling pathways, particularly PI3K/Akt/mTORC1, is described by others to underlie their senolytic effects (Di Micco et al. 2021; Tavenier et al. 2024). Indeed, evidence shows that flavonoids like fisetin can suppress these pathways *in vitro* and *in vivo* (Syed et al. 2014; Adhami et al. 2012; Chamcheu et al. 2019; Mazurakova et al. 2024). Nonetheless, our data suggest that non-plant-based antioxidants can also inhibit PI3K/Akt/mTORC1 signalling in senescent cells and, therefore, the PI3K/Akt/mTORC1 inhibition observed with fisetin, might be downstream of its antioxidant properties. Together with the observation that antioxidants alone can selectively eliminate senescent cells, we propose that, *in vitro*, the senolytic properties of several plant-derived compounds may, at least in part, stem from their ability to scavenge ROS.

Our data strongly suggest that manipulating ROS can be exploited to target PI3K/Akt/mTORC1 signalling and that prolonged treatment can result in senescent cell death or alleviation of senescent phenotypes. However, it is important to acknowledge the limitations of this study. First, the model used in this study may not fully reflect the senescent phenotype observed *in vivo*. Most experiments were conducted using the C2C12 mouse myoblast cell line, which, unlike primary myoblasts or myogenic cells *in vivo* (Sousa-Victor et al. 2014), does not appear to express functional p16^INK4a^ (Kamal et al. 2023), a cyclin-dependent kinase inhibitor involved in the acquisition of the senescent phenotype (Di Micco et al. 2021). Furthermore, the capacity of MitoQ to mitigate mitochondrial ROS production and oxidative damage *in vitro* is well-established (Kelso et al. 2001). However, evidence from cancer cell lines has shown that MitoQ can induce cytotoxicity even at low doses through inhibition of complex I of the electron transport chain (Cheng et al. 2023). In the absence of a redox-inactive MitoQ or an analogous negative control, the cytotoxic effects observed in our study should be interpreted with relative caution. Lastly, this study used proliferating cells as the “control”, a choice that carries inherent limitations, particularly when evaluating cell death. In a hypothetical scenario where proliferating and senescent cells undergo cell death at comparable rates, the continuous division of surviving proliferating cells could mask the true extent of cell loss. While this is the standard approach for studying senolytics *in vitro*, to our knowledge this specific limitation has not yet been addressed.

## Materials and Methods

### Reagents and Cell Culture

List of reagents is shown in Table 1. C2C12 murine skeletal myoblasts (ECACC, Sigma-Aldrich) and primary human myoblasts were grown in Growth Medium (GM), consisting of high-glucose Dulbecco’s Modified Eagle’s Medium (DMEM; Sigma-Aldrich or Gibco), supplemented with 20% fetal bovine serum (PAN Biotech, Aidenbach, Germany) and 1% penicillin-streptomycin (Thermo Fisher Scientific).

Human myoblasts were isolated from human skeletal muscle biopsies as previously described (Gagnon et al. 2025), sorted for the presence of myogenic cell surface marker CD56 using MidiMACS™ system (Miltenyi Biotech), and were cultured in dishes pre-coated with 10% gelatin in PBS.

### Senescence induction, starvation and treatment

To induce cell senescence, myoblasts at approximately 50% confluency were treated with 5 and 10 µM of etoposide for 48 h for C2C12 and human myoblasts, respectively, and then grown in GM for additional five days. The starvation protocol was initiated on day 4 of etoposide treatment. Cells were cultured in serum-free DMEM containing 1% PS for 18 h, followed by 1 h of combined serum and amino acid deprivation in glucose-free DMEM (USBiological Life Sciences) supplemented with 1% PS and 4.5 g/L glucose (Sigma-Aldrich). Treatments were applied concomitantly during starvation at the indicated doses.

### Autophagic flux

Autophagic flux was calculated from Western blot data. LC3-II levels were normalised to α-Tubulin, and flux was determined by subtracting LC3-II in vehicle-treated cells from LC3-II in chloroquine-treated cells.

### Protein synthesis

Protein synthesis was measured using the SUnSET method (Schmidt *et al*., 2009). Which relies upon incorporation of the antibiotic and tyrosyl-tRNA analogue puromycin into nascent proteins. Briefly, 1 µM puromycin (Sigma) was added to control or senescent cells for 1 hour during amino acid starvation prior to cell lysis and subsequent immunoblotting.

### Myoblast Differentiation

C2C12 myoblasts were cultured in 96-well plates to 95–100% confluency. Senescent cells were initially grown in T175 flasks and re-plated on day two post-etoposide at high density (81,250 cells/cm^2^) in 96-well plates and maintained for three more days before differentiation. Differentiation was induced by replacing growth medium (GM) with differentiation medium (DM: high-glucose DMEM + 2% horse serum + 1% penicillin-streptomycin) for five days. Drug treatment was applied either 24 h before differentiation (pretreatment) or 24 h before and throughout differentiation (prolonged). DM from pretreated cells was collected on days 3 and 5, centrifuged (1,000 × g, 10 min), and stored at −80 °C. This DM was mixed 1:1 with fresh DM to produce conditioned medium (CM) for inducing differentiation in fresh control cells.

### DCFDA and MitoSOX live imaging

For mitochondrial superoxide production, cells were treated with 0.5 µM of MitoSOX Red co-stained with 10 nM of MitoTracker Deep Red (both Invitrogen) to normalise for total mitochondrial content. After five minutes, the cells were washed three times and imaged in Fluorobrite DMEM media (Gibco). Intracellular oxidants were quantified using 5 µM of fluorogenic dye 2′,7′-dichlorofluorescin diacetate (DCFDA, Sigma Aldrich). After 20 minutes, cells were washed once and imaged using a LEICA DMIL LED inverted microscope with a Leica DFC360 FX camera. Fluorescence intensity was determined using Fiji/ImageJ by obtaining the raw integrated density value per cell. The coefficient of variability was calculated by dividing SD by mean.

### Cell death live imaging

C2C12 myoblasts in 96-well plates were treated with the indicated drugs for 3 days, of which the first 19 h were during the starvation protocol, to ensure that myoblasts won’t reach confluence and differentiate. Cell death was quantified as the percentage of Hoechst 33342-stained cells, a measure of total nuclei, being co-stained with SYTOX (NucGreen™ Dead 488 ReadyProbes™ Reagent), which accumulates in permeabilised nuclei, indicative of cell death. Cells were imaged at the exact centre of each well using a LEICA DMIL LED inverted microscope equipped with a Leica DFC360 FX camera. Nuclei were counted using Fiji/ImageJ particle analysis tool.

### Immunofluorescence

Cells were fixed in 4% formaldehyde in PBS for 10 minutes and then washed three times for 5 minutes in PBS. Permeabilisation and blocking were performed using either a combined solution of 5% goat serum and 0.2% Triton X-100 for 1 hour (differentiation), or by permeabilising with 0.2% Triton X-100 for 10 minutes followed by blocking in 5% goat serum for 1 h (mTOR/LAMP1 colocalization). Cells were then incubated overnight at 4 °C with the following primary antibodies: rabbit anti-MF-20 (1:200, DSHB), rabbit anti-mTOR (1:200, #2983, Cell Signaling Technology), and rat anti-LAMP1 (1:100, ab208943, Abcam). Cells were then washed and stained with DAPI (1:1000, #62248, Thermo Fisher Scientific) and Rhodamine–phalloidin conjugate (1:500, #10063052, Invitrogen) for actin staining, along with appropriate secondary antibodies: goat anti-rabbit Alexa Fluor 488 (1:500, ab15077, Abcam), goat anti-rabbit Alexa Fluor 647 (1:500, #A21245, Invitrogen), and goat anti-rat Alexa Fluor 488 (1:500, #A48262, Invitrogen).

After an hour, cells were washed and mounted on glass slides using Fluoromount mounting medium (F4680, Sigma-Aldrich) and imaged using a Leica DM2500 microscope with a Leica DFC360 FX camera for qualitative assessment of senescent and control cell morphology. For mTOR/LAMP1 colocalization, confocal images were acquired using a Nikon Ti2 microscope with a 100× oil-immersion objective and NIS Elements software. Colocalization was quantified in ImageJ using Manders’ coefficients. For MF-20 and phalloidin-stained cells in 96-well plates, samples were washed and stored in PBS at 4 °C and imaged within three days using a BioTek Cytation 5 Cell Imaging Multimode Reader (Agilent). Differentiation was quantified in the upper three-quarters of each well (to avoid pipetting artefacts in the 96-well plate) by measuring myotube diameter at standardised locations using the “Straight” tool in Fiji/ImageJ, and by measuring MyHC^+^ coverage. For senescent myoblasts, nonspecific staining was excluded using Fiji/ImageJ’s particle analysis tool with size exclusion, and MyHC^+^ coverage was measured.

### β-Galactosidase assay

Senescence β-Galactosidase Staining Kit (9860, Cell Signaling Technology) was used according to the manufacturer’s instructions. Cells were fixed 1× fixative for 10 minutes, washed, and stored at 4 °C until staining. Staining with the final β-Galactosidase Staining Solution was carried out overnight at 37 °C in a dry (non-CO_2_) incubator. The following day, images were acquired at 10× magnification using a Leica DM IL LED microscope with an MC170 HD camera under brightfield or phase contrast. β-Gal–positive cells were manually counted and expressed as a percentage of total cells.

### Immunoblotting

Cells at low confluency in T25 or T75 flasks were lysed in a small volume of RIPA buffer supplemented with Halt™ protease and phosphatase inhibitors (Thermo Fisher Scientific) and mixed thoroughly. Lysates were kept on ice for approximately 1 hour and then centrifuged at 12,000 × *g* for 10 minutes to remove insoluble material. Protein concentration was determined using the Pierce™ BCA Protein Assay Kit (Thermo Fisher Scientific). Lysates were then mixed with 4× Laemmli buffer, β-mercaptoethanol, and distilled water, boiled at 95 °C for 5 minutes, and loaded onto SDS-PAGE gels at equal protein amounts (5–10 µg). After electrophoresis, proteins were transferred onto PVDF membranes for 90 minutes at 0.35 A. Membranes were washed in Tris-buffered saline containing 0.1% Tween-20 (TBST; Sigma-Aldrich), blocked in milk, BSA, or EveryBlot™ Blocking Buffer (Bio-Rad), and probed with antibodies as detailed in Table S2. Membranes were incubated with Clarity™ Western ECL Substrate (Bio-Rad) for 5 minutes in the dark. Signals were detected using a ChemiDoc imaging system (Bio-Rad) and quantified using the lane and band analysis tool in ImageLab software. Images were further adjusted in ImageLab for optimal presentation without affecting band quantification.

### RNA extraction and RT-qPCR

RNA was extracted using the TRIzol™ method (Invitrogen) according to the manufacturer’s instructions and dissolved in Ambion™ RNA Storage Solution (Invitrogen). RNA concentration and purity were assessed using UV spectroscopy with a NanoDrop™ 2000 spectrophotometer (Thermo Fisher Scientific). One-step RT-qPCR was performed on a ViiA™ 7 Real-Time PCR System (Applied Biosystems) using a 10 µL reaction volume containing 5 µL of SYBR Green-based one-step PrecisionPLUS qPCR Master Mix, 4.5 µL of diluted RNA (10–25 ng), and 0.5 µL of diluted forward and reverse primers. mRNA expression was calculated using the ΔΔCT method and normalised to GAPDH or RP2. The primer sequences are shown in Table S3.

### Statistical Analysis

Results are reported as mean ± standard deviation in either absolute values or normalised using fixed point or sum normalisation for comparison between two or more conditions, respectively. Statistical analyses were performed using heteroscedastic *t*-tests in Microsoft Excel or with the ggpubr, rstatix, EnvStats, and DescTools packages in RStudio. The specific statistical tests used are indicated in the corresponding figure legends.

## Acknowledgements

The authors are grateful to Antonios Giannopoulos and José Rui Rodrigues for their assistance with handling human cells. The authors would like to thank Dr Ryan Brett for his assistance with confocal microscopy.

## Authors’ contributions

V.B. and N.R.W.M. conceptualised the research project and designed the experiments. V.B., A.D., B.T., M.T., S.D.G., J.Q., H.W., E.W., N.M. and N.R.W.M. performed experiments. V.B. and E.C. analysed experimental data. V.B. and N.R.W.M. wrote the manuscript. V.B., O.G.D., H.F.D., M.T., B.C. and N.R.W.M. contributed to the methodology and helped revise the manuscript. V.B. and N.R.W.M. acquired funding. All authors have read and agreed to the published version of the manuscript.

## Data availability

The data that support the findings of this study are available from the corresponding author upon reasonable request.

