## Supplementary Material for "Senescent myoblasts exhibit ROS-dependent Akt-mTORC1 dysregulation and are susceptible to reductive stress-induced cell death"

### Supplementary Figures

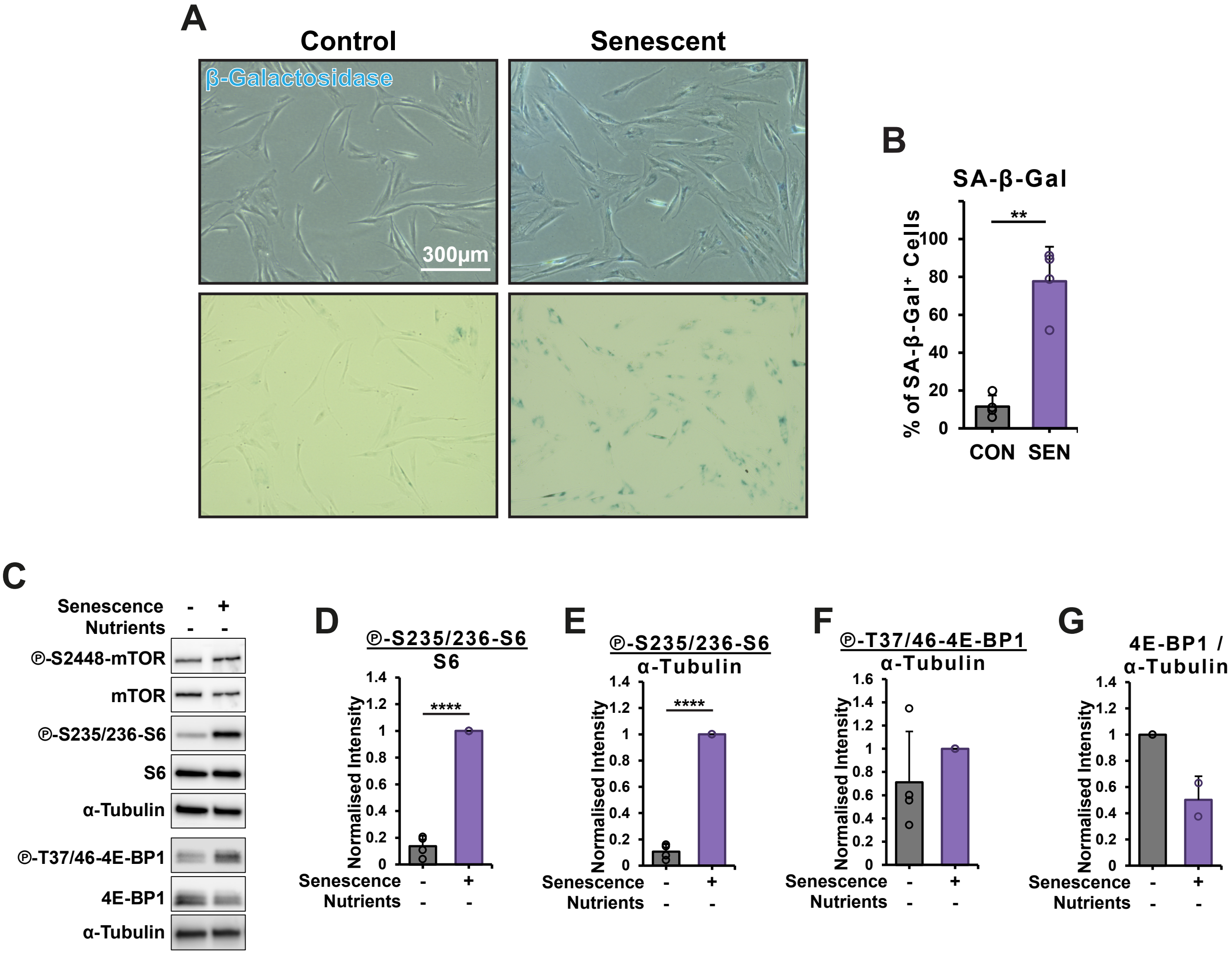


**Figure S1. mTORC1 dysregulation in primary human myoblasts.**

Primary human myoblasts were treated with 10 μM etoposide for 48 h and then maintained in growth medium for an additional five days (A and B) and starved in the last 19 hours (C-G).

(A and B) β-galactosidase staining and quantification.

(C-G) Immunoblot analysis with quantification of mTORC1 signalling markers

Results are reported as mean ± standard deviation. P values were calculated using independent, heteroscedastic t-test. n = 4, except for G, where n = 2. $^{**}P \leq 0.01$, $^{****}P \leq 0.0001$. CON = control, SEN = senescent.

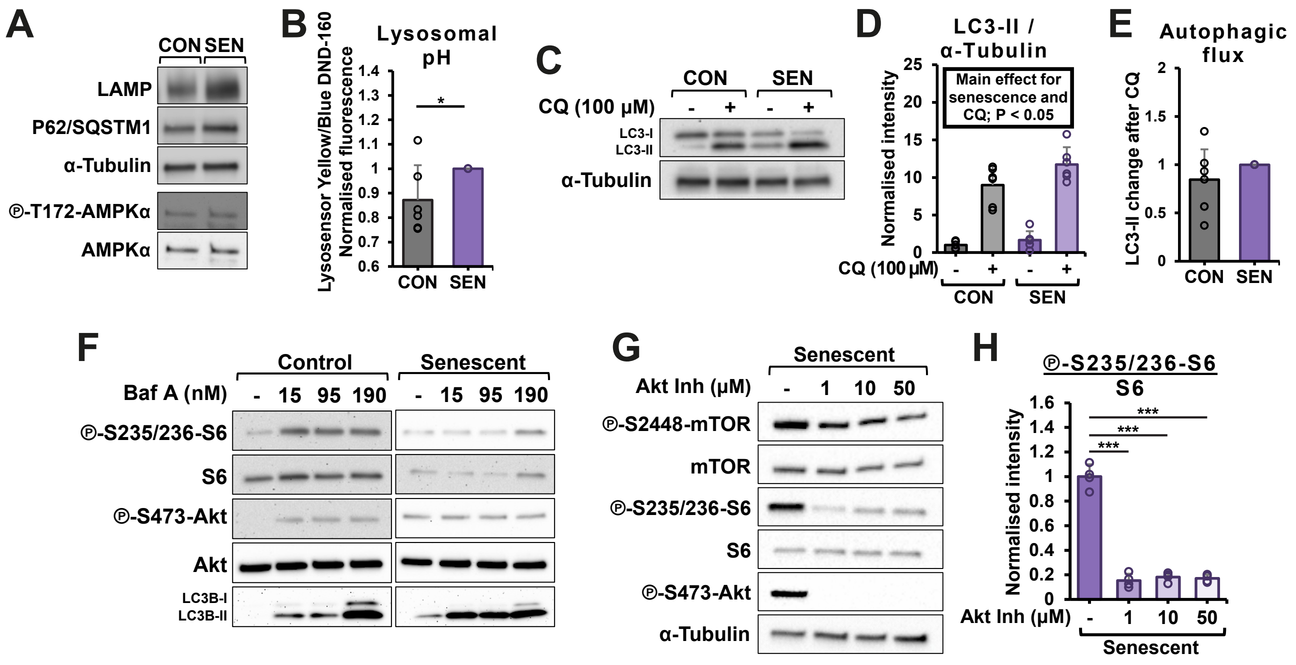


**Figure S2. Inhibition of Akt, but not autophagy, alleviates mTORC1 activity in starved senescent C2C12 myoblasts.**

(A, C, D and F-H) Immunoblot analysis and quantification for Akt/mTORC1 and autophagy-related proteins in starved senescent C2C12 myoblasts treated with the indicated compound during the last hour of standard starvation protocol.

(E) Autophagic flux calculation based on LC3-II expression.

(B) Lysosomal pH in non-starved control and senescent myoblasts, measured using Lysosensor Yellow/Blue DND-160.

P values were calculated using an independent, heteroscedastic t-test (B and E), one-way ANOVA, (D and H) followed by Dunnett's post hoc test (H). n = 2 for F and 3–6 for the rest. *$P \leq 0.05$, **$P \leq 0.01$, ***$P \leq 0.001$, ****$P \leq 0.0001$. CQ = chloroquine, Baf A = Bafilomycin A1, Akt Inh = Akt inhibitor, CON = control, SEN = senescent.

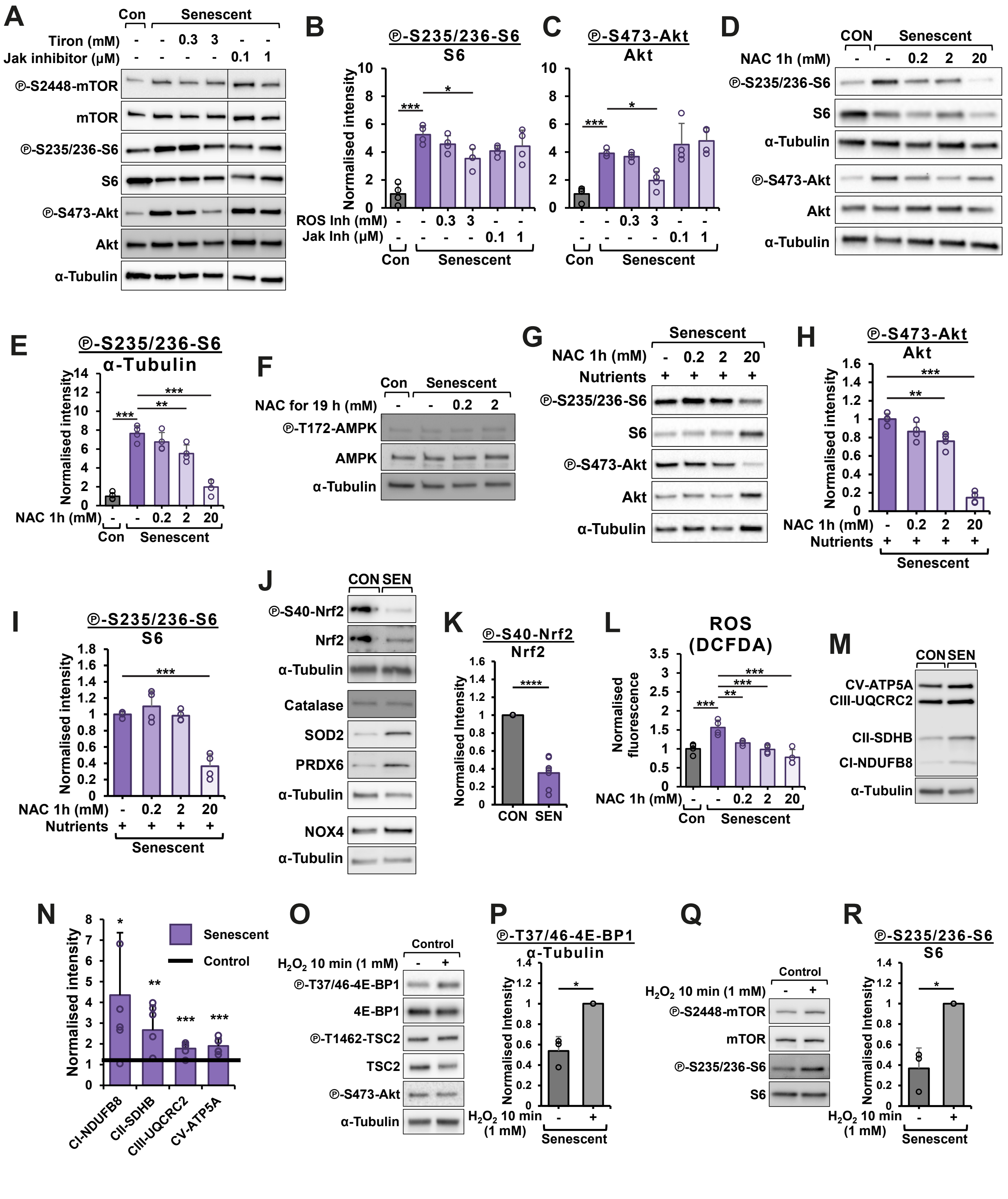
A
B
C
D
E
F
G
H
I
J
K
L
M
N
O
P
Q
R
Ⓟ-S2448-mTOR
mTOR
Ⓟ-S235/236-S6
S6
Ⓟ-S473-Akt
Akt
α-Tubulin
Tiron (mM)
Jak inhibitor (µM)
Con
Senescent
Normalised intensity
ROS Inh (mM)
Jak Inh (µM)
NAC 1h (mM)
CON
NAC for 19 h (mM)
Ⓟ-T172-AMPK
AMPK
Nutrients
Ⓟ-S40-Nrf2
Nrf2
Catalase
SOD2
PRDX6
NOX4
SEN
ROS
(DCFDA)
Normalised fluorescence
CV-ATP5A
CIII-UQCRC2
CII-SDHB
CI-NDUFB8
Control
$H_2O_2$ 10 min (1 mM)
Ⓟ-T37/46-4E-BP1
4E-BP1
Ⓟ-T1462-TSC2
TSC2

**Figure S3. Interaction between Akt/mTORC1 signalling and ROS.**

(A-C) Western blot analysis for Akt/mTORC1 signalling in starved senescent C2C12 myoblasts treated with the indicated doses of ROS scavenger/antioxidant tiron and Jak inhibitor tofacitinib.

(D and E) Western blot analysis for Akt/mTORC1 signalling in starved senescent C2C12 myoblasts kept treated with indicated doses of NAC for one hour.

(F) Western blot analysis for AMPK signalling in starved senescent C2C12 myoblasts treated with the indicated doses of NAC for 19 hours.

(G-I) Immunoblot analysis and quantification for Akt/mTORC1 signalling in senescent C2C12 myoblasts kept in full GM treated with indicated doses of NAC for one hour.

(J and K) Western blot analysis of the transcriptional regulator of the antioxidant response (Nrf2, K), endogenous antioxidants (catalase, SOD2, PRDX6), and NOX4 in starved myoblasts.

(L) DCFDA ROS assay quantified using a microplate reader in myoblasts treated with the indicated doses of NAC for 1 h.

(M and N) Western blot analysis of ETC complexes in starved control and senescent cells.

(O–R) Western blot analysis of Akt/mTORC1 signalling in starved control cells treated with $H_2O_2$.

Results are reported as mean ± standard deviation. P values were calculated using an independent, heteroscedastic t-test (K, N, P, R) and one way ANOVA followed by Dunnett's post hoc test (B, C, E, H, I, L). n = 3-7. $^*P \leq 0.05$, $^{**}P \leq 0.01$, $^{***}P \leq 0.001$, $^{****}P \leq 0.0001$. ROS Inh = ROS inhibitor/antioxidant Tiron, Jak Inh = Jak inhibitor Tofacitinib, NAC = N-acetylcysteine, Con = control cells

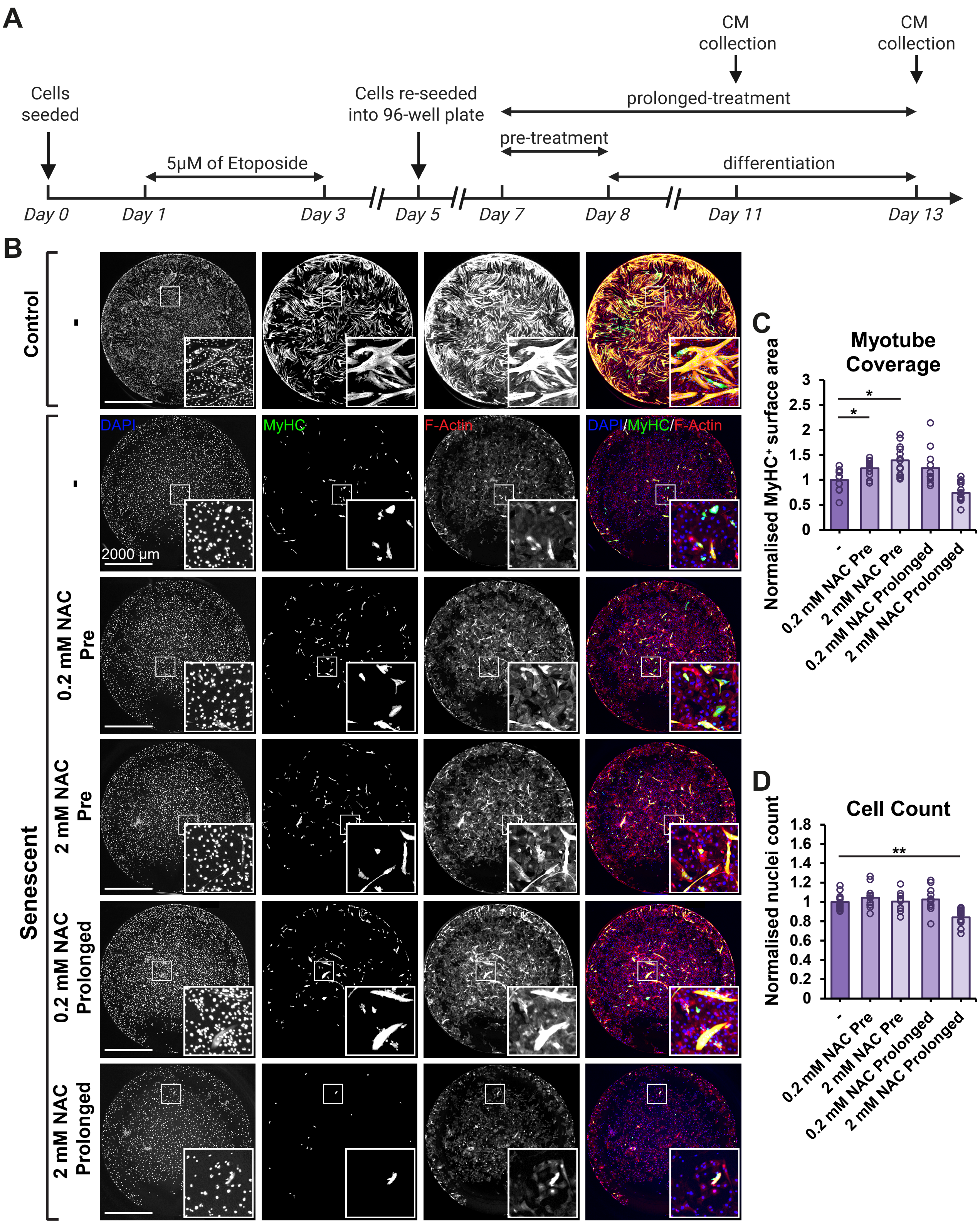
A
CM collection
CM collection
Cells seeded
Cells re-seeded into 96-well plate
prolonged-treatment
pre-treatment
5µM of Etoposide
differentiation
Day 0
Day 1
Day 3
Day 5
Day 7
Day 8
Day 11
Day 13
B
Control
Senescent
-
0.2 mM NAC Pre
2 mM NAC Pre
0.2 mM NAC Prolonged
2 mM NAC Prolonged
DAPI
MyHC
F-Actin
DAPI/MyHC/F-Actin
2000 µm
C
Myotube Coverage
Normalised MyHC⁺ surface area
0
0.5
1
1.5
2
2.5
3
*
*
-
0.2 mM NAC Pre
2 mM NAC Pre
0.2 mM NAC Prolonged
2 mM NAC Prolonged
D
Cell Count
Normalised nuclei count
0
0.2
0.4
0.6
0.8
1
1.2
1.4
1.6
1.8
**
-
0.2 mM NAC Pre
2 mM NAC Pre
0.2 mM NAC Prolonged
2 mM NAC Prolonged

**Figure S4. The effect of antioxidants on myotube differentiation in senescent C2C12 myoblasts**

(A) Protocol summary for all differentiation experiments.

(B-D) Representative images (B) of differentiated senescent C2C12 myoblasts stained for F-Actin and MyHC, and quantification of myotube coverage (C) and cell count (DAPI staining, D). Cells were stimulated to differentiate after 24 h pretreatment (Pre) or after 24 h pretreatment combined with concomitant antioxidant treatment during differentiation (Prolonged).

Results are reported as mean ± standard deviation. P values were calculated using one way ANOVA followed by Dunnett's post hoc test. n = 9-12. *P ≤ 0.05, **P ≤ 0.01. NAC = N-acetylcysteine, MitoQ = Mitoquinone mesylate, CON = control, SEN = senescent.

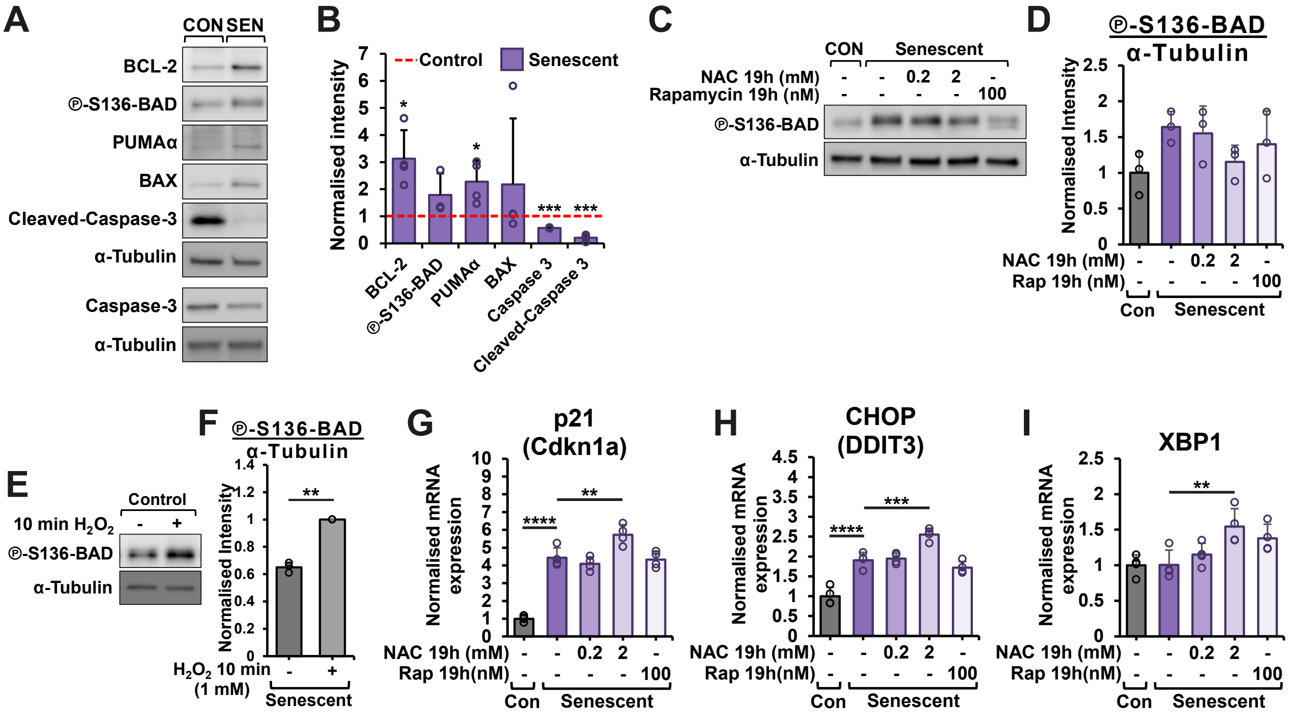


**Figure S5. Senescent C2C12 myoblasts are susceptible to antioxidant-induced cell death.**

(A and B) Western blot analysis for cell survival related signalling in control and senescent myoblasts.

(C-F) Western blot analysis of for phosphorylation of BAD in control and senescent myoblasts treated with NAC, rapamycin and H2O2 as indicated.

(G-I) mRNA expression of DNA damage and ER stress-related genes.

Results are reported as mean ± standard deviation. P values were calculated using an independent, heteroscedastic t-test (F), and one-way ANOVA followed by Dunnett's post hoc test (the rest). *P ≤ 0.05, **P ≤ 0.01, ***P ≤ 0.001, ****P≤ 0.0001. CON = control, SEN = senescent, NAC = N-acetylcysteine, Rap = rapamycin.

### Supplementary Methods

Lysosenor and DCFDA plate-reader based assays

Lysosensor Yellow/Blue DND-160 (Thermo Fisher Scientific) was used to measure lysosomal pH. Cells were treated with 5 µM Lysosensor in PBS for 20 minutes, followed by three PBS washes. Fluorescence readings were taken from the bottom of the plate using a Varioskan™ LUX plate reader (Thermo Fisher Scientific) every minute for 10 minutes. To assess cellular oxidant production, cells were treated with 5 µM DCFDA for 1 hour concomitantly with the drug treatment, and fluorescence was subsequently measured using the microplate reader.

To account for differences in cell size and proliferation rates between control and senescent cells during plate-reader measurements, fluorescence values were normalised to protein concentration. Control cells were seeded into black 96-well plates at varying densities, while senescent cells were grown in parallel on the same plate. Following fluorescence acquisition, cells were lysed using RIPA buffer, and protein concentration was determined using the Pierce™ BCA Protein Assay Kit (Thermo Fisher Scientific). A regression analysis was performed using protein concentrations from control cells (y-axis) against their corresponding fluorescence values (x-axis). The protein concentration of senescent cells was then used to estimate an equivalent fluorescence value for control cells, enabling direct comparison between control and senescent samples.

### Supplementary Tables

#### Table S1. Reagents

| Cell culture chemicals | Source | Identifier |
|---|---|---|
| C2C12 | ECACC | 91031101 |
| DMEM | Sigma-Aldrich | D6429 |
| Etoposide | Sigma-Aldrich | 341205 |
| FluoroBrite DMEM | Gibco | A1896701 |
| DMEM w/o Amino Acids, Glucose (Powder) | USBiological | D9800-27 |
| MEM Amino Acid [50X] | Gibco | 11130051 |
| D-(+)-Glucose | Sigma-Aldrich | G7021 |
| Sodium Bicarbonate | Sigma-Aldrich | S5761 |
| Horse serum | Sigma-Aldrich | H1270 |
| Pen Strep | Gibco | 15140122 |
| Trypsin-EDTA | Gibco | 11590626 |
| StemPro Accutase | Gibco | A1110501 |
| Gelatin solution | Sigma-Aldrich by Merck | G1393 |
| N-Acetyl-LCysteine | Sigma-Aldrich by Merck | A9165 |
| Tiron | Thermo Fisher Scientific | 174140250 |

| | | |
|---|---|---|
| MitoQ (Mitoquinone mesylate) | MedChemExpress | HY-100116A |
| Mito-TEMPO | MedChemExpress | HY-112879 |
| GLX351322 | MedChemExpress | HY-100111 |
| GKT137831 | Cayman chemical | 17764 |
| DPI (Diphenyleneiodonium chloride) | Sigma-Aldrich | D2926 |
| Rotenone | Sigma-Aldrich | R8875 |
| Hydrogen peroxide solution | Sigma-Aldrich | H1009 |
| TBHP | Abcam | ab113851 |
| Sodium phenylbutyrate (4PBA) | Sigma-Aldrich | SML0309 |
| Tofacitinib (CP-690550) Citrate | APExBIO | A4135 |
| BMS-345541 | APExBIO | B4655 |
| Chloroquine diphosphate salt | Sigma-Aldrich | C6628 |
| Bafilomycin A1 | Sigma-Aldrich | SML1661 |
| Akt Inhibitor | Sigma-Aldrich | 124005 |
| Rapamycin | Sigma-Aldrich | 553211 |
| Nigericin sodium salt | Sigma-Aldrich | N7143 |
| Monensin sodium salt | Sigma-Aldrich | M5273 |
| TRIzol | Invitrogen | 15596026 |
| Pierce RIPA buffer | Thermo Fisher Scientific | 89900 |
| Halt Protease and Phosphatase inhibitor Cocktail | Thermo Fisher Scientific | 78440 |
| **Commercial assay kits** | **Source** | **Identifier** |
| Pierce BCA Protein Assay Kit | Thermo Fisher Scientific | 23227 |
| β-Galactosidase staining | Cell Signalling Technology | #9860 |
| **Fluorescent dyes** | **Source** | **Identifier** |
| LysoSensor Yellow/Blue DND-160 | Invitrogen | L7545 |
| DCFDA | Sigma-Aldrich | 287810 |
| MitoSOX Red | Invitrogen | M36008 |
| MitoTracker Deep Red FM | Invitrogen | M22426 |
| MitoTracker Orange CMTMRos | Invitrogen | M7510 |
| DAPI | Thermo Fisher Scientific | 62248 |
| Hoechst 33342 | Thermo Fisher Scientific | 62249 |
| NucGreen™ Dead 488 (SYTOX™ Green) | Invitrogen | R37109 |
| **Antibodies** | **Source** | **Identifier** |
| ℗-S1101-IRS-1 | Cell Signalling Technology | #2385 |
| ℗-T1462-TSC2 | Cell Signalling Technology | #3611 |
| Tuberin/TSC2 | Cell Signalling Technology | #4308 |
| ℗-S2448-mTOR | Cell Signalling Technology | #5536 |
| mTOR | Cell Signalling Technology | #2983 |
| ℗-T389-S6K1 | Cell Signalling Technology | #9234 |

| | | |
|---|---|---|
| S6K1 | Cell Signalling Technology | #2708 |
| ℗-S235/236-S6 | Cell Signalling Technology | #2211 |
| S6 | Cell Signalling Technology | #2217 |
| α-Tubulin | Cell Signalling Technology | #2125 |
| ℗-T37/46-4E-BP1 | Cell Signalling Technology | #2855 |
| 4E-BP1 | Cell Signalling Technology | #2855 |
| ℗-T458/199 PI3K p85/p55 | Cell Signalling Technology | #4228 |
| PI3K p85 | Cell Signalling Technology | #4292 |
| ℗-S473-Akt | Cell Signalling Technology | #4060 |
| Akt | Cell Signalling Technology | #4691 |
| ℗-T172-AMPK | Cell Signalling Technology | #2531 |
| AMPK | Cell Signalling Technology | #2532 |
| LAMP1 (immunoblotting) | Cell Signalling Technology | #3243 |
| p62/SQSTM1 | Cell Signalling Technology | #5114 |
| LC3 | Cell Signalling Technology | #3868 |
| NOX4 | Proteintech | #14347-1-AP |
| ℗-S40-Nrf2 | Abcam | ab180844 |
| Nrf2 | Abcam | ab62352 |
| Catalase | Cell Signalling Technology | #314097 |
| SOD2 | Cell Signalling Technology | #13141 |
| GSTM3 | Proteintech | #15214-1-AP |
| Total OXPHOS | Abcam | ab110411 |
| PGC-1α | Cell Signalling Technology | #2178 |
| Caspase-3 | Cell Signalling Technology | #14220 |
| Cleaved Caspase-3 | Cell Signalling Technology | #9664 |
| ℗-S136-BAD | Cell Signalling Technology | #4366 |
| BAD | Santa Cruz | sc-8044 |
| BCL-2 | Santa Cruz | sc-7382 |
| BAX | Santa Cruz | sc-7480 |
| PUMAα | Santa Cruz | sc-377015 |
| Anti-Rabbit IgG, HRP-linked | Cell Signalling Technology | #7074 |
| Anti-mouse IgG, HRP-linked | Cell Signalling Technology | #7076 |
| Desmin | Dako | IS606 |
| MF-20 | DSHB | - |
| Rhodamine Phalloidin | Invitrogen | #10063052 |
| LAMP1 (immunocytochemistry) | Abcam | ab208943 |
| Goat anti-mouse Alexa Fluor 488 | Invitrogen | A32723 |
| Goat anti-rat Alexa Fluor 488 | Invitrogen | A48262 |
| Goat anti-rabbit Alexa Fluor 488 | Abcam | ab15077 |

| | | |
|---|---|---|
| Goat anti-rabbit Alexa Fluor 647 | Invitrogen | A21245 |
| Fluoromount$^{TM}$ Aqueous Mounting Medium | Sigma Aldrich | F4680 |

ECACC – European Collection of Authenticated Cell Cultures

**Table S2. Antibody dilutions used for Western blotting.**

| Target | 1ºAb Product No. | Blocking | 1º Ab solution | 2º Ab solution |
|---|---|---|---|---|
| **℗-T1462-TSC2** | CST #3611 | 5% BSA | 1:1000 in 2% milk | 1:5000 in 2 % milk |
| **Tuberin/TSC2** | CST #4308 | 5% BSA | 1:1000 in 2% milk | 1:5000 in 2 % milk |
| **℗-T458/199 PI3K p85/p55** | CST #4228 | 5% BSA | 1:1000 in 2% milk | 1:5000 in 2 % milk |
| **PI3K p85** | CST #4292 | 5% BSA | 1:1000 in 2% milk | 1:5000 in 2 % milk |
| **℗-S2448-mTOR** | CST #5536 | 5% BSA | 1:2000 in 2% BSA | 1:5000 in 2 % BSA |
| **mTOR** | CST #2983 | 5% BSA | 1:2000 in 2% BSA | 1:5000 in 2 % BSA |
| **℗-T389-S6K1** | CST #9234 | 5% milk | 1:500 in 2% milk | 1:5000 in 2 % milk |
| **S6K1** | CST #2708 | 5% milk | 1:500 in 2% milk | 1:5000 in 2 % milk |
| **℗-S235/236-S6** | CST #2211 | 5% milk | 1:5000 in 2% milk | 1:5000 in 2 % milk |
| **S6** | CST #2217 | 5% milk | 1:5000 in 2% milk | 1:5000 in 2 % milk |
| **α-Tubulin** | CST #2125 | 5% milk | 1:2000 in 2% milk | 1:5000 in 2 % milk |
| **℗-T37/46-4E-BP1** | CST #2855 | 5% milk | 1:5000 in 2% milk | 1:5000 in 2 % milk |
| **4E-BP1** | CST #2855 | 5% milk | 1:5000 in 2% milk | 1:5000 in 2 % milk |
| **℗-S473-Akt** | CST #4060 | 5% milk | 1:750 in 2% milk | 1:5000 in 2 % milk |
| **Akt** | CST #4691 | 5% milk | 1:2000 in 2% milk | 1:5000 in 2 % milk |
| **℗-T172-AMPK** | CST #2531 | 5% milk | 1:1000 in 2% milk | 1:5000 in 2 % milk |
| **AMPK** | CST #2532 | 5% milk | 1:1000 in 2% milk | 1:5000 in 2 % milk |
| **LAMP1** | CST #3243 | 5% milk | 1:2000 in 2% in milk | 1:5000 in 2 % milk |
| **p62** | CST #5114 | 5% BSA | 1:1000 in 2% BSA | 1:5000 in 2 % BSA |
| **LC3** | CST #3868 | 5% milk | 1:500 in 2% milk | 1:2000 in 2 % milk |
| **NOX4** | Proteintech #14347-1-AP | 5% milk | 1:1000 in 5% milk | 1:5000 in 5 % milk |
| **℗-S40-Nrf2** | Abcam ab180844 | 5% BSA | 1:2000 in 5% BSA | 1:5000 in 5 % BSA |
| **Nrf2** | Abcam ab62352 | 5% BSA | 1:2000 in 5% BSA | 1:5000 in 5 % BSA |
| **Catalase** | CST #314097 | EveryBlot | 1:1000 in EveryBlot | 1:5000 in EveryBlot |
| **SOD2** | CST #13141 | EveryBlot | 1:5000 in EveryBlot | 1:5000 in EveryBlot |
| **GSTM3** | Proteintech #15214-1-AP | EveryBlot | 1:500 in EveryBlot | 1:5000 in EveryBlot |
| **Total OXPHOS** | Abcam ab110411 | EveryBlot | 1:1000 in EveryBlot | 1:5000 in EveryBlot |
| **Caspase-3** | CST #14220 | 5% milk | 1:500 in 5% BSA | 1:5000 in 5% BSA |
| **Cleaved Caspase-3** | CST #9664 | 5% milk | 1:500 in 5% milk | 1:5000 in 5% BSA |
| **℗-S136-BAD** | CST #4366 | 5% milk | 1:500 in 5% BSA | 1:5000 in 5% BSA |
| **BCL-2** | Santa Cruz sc-7382 | 5% milk | 1:1000 in 5% BSA | 1:5000 in 5% BSA |
| **Bax** | Santa Cruz sc-7480 | 5% milk | 1:500 in 5% BSA | 1:5000 in 5% BSA |
| **PUMAα** | Santa Cruz sc-377015 | 5% milk | 1:500 in 5% BSA | 1:5000 in 5% BSA |

Secondary antibody: anti-mouse IgG, HRP-linked antibody (Cell Signalling Technology, #7076) was used for Total OXHPHOS, BCL-2, and BAX antibodies, the remaining antibodies were bound to anti-rabbit IgG HRP-conjugated (Cell Signalling Technology, #7074). CST = Cell Signalling Technology; BSA = Bovine Serum Albumin.

### Table S3. Primer sequences used for RT-qPCR

| Target Gene (protein) | Primer Sequence (5`-3`) | Reference number |
|---|---|---|
| *Polr2b* (RP2) | F: GGTCAGCCGGGAACTTGTGGTAT<br>R: GCATCATTAAATGGAGTAGCGTC | NM_153798.2 |
| *Gapdh* (GAPDH) | F: TGTGTCCGTCGTGGATCTGA<br>R: TTGCTGTTGAAGTCGCAGGAG | NM_001411843 |
| *Trp53* (p53) | F: TGAAGCACCGCTTCCCGAAGAG<br>R: AGAAGACGACTGGGGCAGCTAT | NM_001007581 |
| *Cdkn1a* (p21) | F: TCGCTGTCTTGCACTCTGGTGT<br>R: CCAATCTGCGCTTGGAGTGATAG | NM_007669 |
| *Cdkn2a* ($p16^{INK4a}$) | F: CGCAGGTTCTTGGTCACTGT<br>R: TGTTCACGAAAGCCAGAGCG | NM_0013283 |
| *Il6* (IL-6) | F: GGTCTGTTGGGAGTGGTATC<br>R: TCCATCCAGTTGCCTTCTTG | NM_031168 |
| *Ccl2* (MCP-1) | F: CAAGATGATCCCAATGAGT<br>R: TTGGTGACAAAAACTACAGC | NM_00178954 |
| *Xbp1s* (XBP1s) | F: GGGCCTGCACCTGCT<br>R: GGGCCTGCACCTGCT | NM_00127173 |
| *Ddit3* (CHOP) | F: CCTCGCTCTCCAGATTCCA<br>R: CTGTTTCCGTTTCCTAGTTCTTC | NM_007837 |

Xbp1 - X-box binding protein 1 spliced, Polr2b – RNA polymerase II polypeptide B, CDKN1A: cyclin-dependent kinase inhibitor 1a, Ddit3: DNA-damage inducible transcript 3, Ccl2: C-C motif chemokine ligand 2, MCP-1: Monocyte Chemoattractant Protein-1, IL-6: interleukin 6; TNF-α tumour necrosis factor-alpha, GAPDH: Glyceraldehyde-3-Phosphate Dehydrogenase.
